# Sustained virome diversity in Antarctic penguins and their ticks: geographical connectedness and no evidence for low pathogen pressure

**DOI:** 10.1101/2019.12.11.873513

**Authors:** Michelle Wille, Erin Harvey, Mang Shi, Daniel Gonzalez-Acuña, Edward C. Holmes, Aeron C. Hurt

**Author notes:** These authors contributed equally to this study (Michelle Wille, Erin Harvey). Corresponding authors: Michelle Wille - Edward C. Holmes.

## Abstract

Despite its isolation and extreme climate, Antarctica is home to diverse fauna and associated microorganisms. It has been proposed that the most iconic Antarctic animal, the penguin, experiences low pathogen pressure, accounting for their disease susceptibility in foreign environments. However, there is a limited understanding of virome diversity in Antarctic species, the extent of *in situ* virus evolution, or how it relates to that in other geographic regions. To test the idea that penguins have limited microbial diversity we determined the viromes of three species of penguins and their ticks sampled on the Antarctic peninsula. Using total RNA-Sequencing we identified 107 viral species, comprising likely penguin associated viruses (n = 13), penguin diet and microbiome associated viruses (n = 82) and tick viruses (n = 8), two of which may have the potential to infect penguins. Notably, the level of virome diversity revealed in penguins is comparable to that seen in Australian waterbirds, including many of the same viral families. These data therefore reject the theory that penguins are subject to lower pathogen pressure. The repeated detection of specific viruses in Antarctic penguins also suggests that rather than being simply spill-over hosts, these animals may act as key virus reservoirs.

## Introduction

Geographical separation and extreme climates have resulted in the long isolation of Antarctica and the subantarctic islands. The result is a unique assemblage of animals, some relying entirely on the frozen continent, with others utilizing the fringes. Such geographic isolation has been proposed to explain why Antarctic fauna supposedly harbour a paucity of viruses, and supported by the observation that captive Antarctic penguins are highly susceptible to infectious diseases (1). It has therefore been hypothesized that Antarctic fauna have evolved in a setting of low “pathogen pressure”, reflected in limited microbial diversity and abundance (1, 2). As a consequence, the potential for climate driven and human mediated movement of microorganisms makes the expansion of infectious diseases to the Antarctic a matter of considerable concern (1, 3–5).

To date, a small number of viral species have been described in Antarctic fauna (6). Serological studies have revealed that Antarctic penguins are reservoirs for influenza A virus (IAV), avian avulaviruses (formerly avian paramyxoviruses), birnaviruses, herpesviruses, and flaviviruses (7–13). Despite improvements in the molecular tools for virus detection, it is only in recent years that full viral genomes have been characterized (6). For example, adenoviruses, astroviruses, paramyxoviruses, orthomyxoviruses, polyomaviruses and papillomavirus have been identified in Adélie penguins (*Pygoscelis adeliae*), Chinstrap penguins (*Pygoscelis antarctica*) and Gentoo penguins (*Pygoscelis papua*) (6, 14–22). However, sampling is limited and genomic data sparse, such that we have a fragmented understanding of virus diversity in penguins and in Antarctica in general.

Antarctic penguins may also be infected by viruses spread by ectoparasites, particularly the seabird tick *Ixodes uriae* (White) (23). For example, seven different arthropod-borne viruses (arboviruses) were identified in *I. uriae* ticks collected from King penguin colonies on Macquarie Island in the subantarctic (25): Nugget virus, Taggert virus, Fish Creek virus, Precarious Point virus, and Catch me-cave virus, all of which belong to the order *Bunyavirales* (*Nairoviridae* and *Phleboviridae*), a member of the *Reoviridae* (Sandy Bay virus, genus *Orbivurus*), and a member of the *Flaviviridae* (Gadgets Gulls virus) (23–25). Notably, *I. uriae* is the only species of tick with a circumpolar distribution and is found across the Antarctic peninsula (26, 27). *I. uriae* are mainly associated with nesting seabirds and feed on penguins in the summer months, using off-host aggregation sites for the reminder of the year (28–30).

Despite concern over virus emergence in Antarctica there remains little understanding of virus diversity in Antarctic species, nor how virome diversity in Antarctic species relates to that seen in other geographic regions. Herein, we determined the viromes of three species of penguins (Gentoo, Chinstrap and Adelie) on the Antarctic peninsula, as well as *I. uriae* ticks that parasitise these birds. With these data in hand we addressed the drivers of virus ecology and evolution in this remote and unique locality.

## Methods

### Ethics statement

Approvals to conduct sampling in Antarctica were provided by the Universidad de Concepción, Facultad de Ciencias Veterinarias, Chillán, Chile (application CBE-48-2013), and the Instituto Antártico Chileno, Chile (application 654).

### Sample collection

Samples were collected as previously described (15, 16). Briefly, samples were collected from the South Shetland Islands and the Antarctic peninsula from Gentoo, Chinstrap, and Adelie penguins in 2014, 2015 and 2016, respectively (Table S1). A cloacal swab was collected from each penguin a using a sterile-tipped swab, placed in viral transport media, and stored at −80 °C within 4-8 hours of collection.

Gentoo, Chinstrap and Adelie penguins were sampled on Kopaitik Island, Rada Covadonga, 1 km west of Base General Bernardo O’Higgins (63°19’5”S, 57°53’55”W). Kopaitik Island is a mixed colony containing these three penguin species, although no survey has been performed since 1996 (31). Gentoo penguins were also sampled adjacent to González Videla Base, Paradise Bay (64°49′26″S 62°51′25″W): this colony is comprised of almost only Gentoo penguins (3915 nests) and a single Chinstrap penguin nest reported in 2017 (31). Chinstrap penguins were sampled at Punta Narebski, King George Island (62°14′00″S 58°46′00″W) and Adelie penguins were sampled at Arctowski Station, Admiralty Bay, King George Island (62°9’35”S, 58°28’17”W). A census at the penguin colony at Punta Narebski in 2013 reported 3157 Chinstrap and 2378 Gentoo penguin nests. The penguin colony adjacent to Arctowski Station comprises both Adelie and Chinstrap penguins, with a 2013 census reporting 3246 and 6123 nests and 3627 and 6595 chicks, respectively (31) (Fig 1). In 2017, the common seabird tick *I. uriae* at various life stages (adult male, adult female and nymphs) were collected from Paradise Bay (Fig 1). Ticks were collected under rocks within and directly adjacent to penguin colonies and placed in RNAlater (Ambion) and stored at −80°C within 4-8 hours of collection.

**Figure 1.**
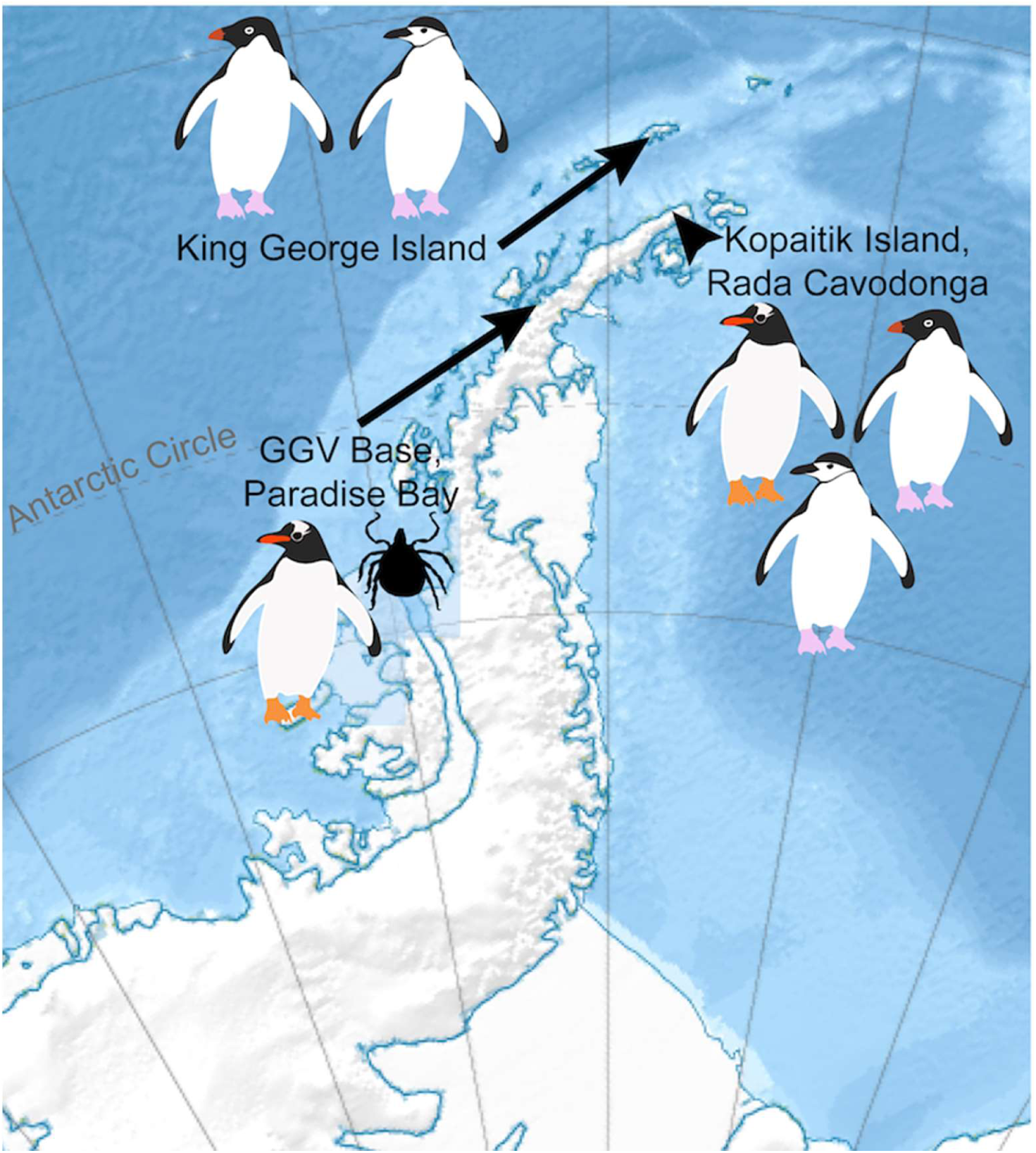
Map of Antarctic peninsula and locations where Antarctic penguins samples and ticks were collected.

### RNA library construction and sequencing

RNA library construction, sequencing and RNA virus discovery was carried out as described previously (32). Briefly, cloacal swab samples from penguins were extracted using the MagMax *mir*Vana™ Total RNA isolation Kit (ThermoFisher Scientific) on the KingFisher™ Flex Purification System (ThermoFisher Scientific). Extracted samples were assessed for RNA quality using the TapeStation 2200 and High Sensitivity RNA reagents (Aligent Genomics, Integrated Sciences).

RNA was extracted from ticks as described previously (33). Briefly, ticks were washed in ethanol and homogenised in lysis buffer using a TissueRuptor (Qiagen) and RNA was extracted using the RNeasy plus mini kit (Qiagen). The quality and concentration of extracted RNA was assessed using the Agilent 4200 TapeStation. The 10 penguin and 5 tick samples with the highest concentration corresponding to species/location/age were then pooled using equal concentrations and concentrated using the RNeasy MinElute Cleanup Kit (Qiagen) (Table S1).

Libraries were constructed using the TruSeq total RNA library preparation protocol (Illumina) and host rRNA was removed using the Ribo-Zero-Gold kit (Illumina) for penguin libraries and the Ribo-Zero Gold rRNA Removal (Epidemiology) kit (Illumina) for the tick libraries. Paired-end sequencing (100bp) of the RNA library was performed on the Illumina HiSeq 2500 platform at the Australian Genome Research Facility (Melbourne). All sequence reads have been deposited in the Sequence Read Archive (XXXX). Virus consensus sequences have been deposited on GenBank (XXXX-XXX)

### RNA virus discovery

Sequence reads were demultiplexed and trimmed with Trimmomatic followed by de novo assembly using Trinity (34). No filtering of host/bacterial reads was performed before assembly. All assembled contigs were compared to the entire non-redundant nucleotide (nt) and protein (nr) database using blastn and diamond blast (35), respectively, setting an e-value threshold of 1×10^−10^ to remove false-positives.

Abundance estimates for all contigs were determined using the RSEM algorithm (34). All contigs that returned blast hits with paired abundance estimates were filtered to remove plant, invertebrate fungal, bacterial and host sequences. Viruses detected in the penguin libraries were divided into those likely to infect birds and those likely associated other hosts (36)(37). This division was performed using a combination of phylogenetic analysis and information on virus associations available at the Virus-Host database (http://www.genome.jp/virushostdb/). The list was cross-referenced with known laboratory reagent contaminants (38). Novel viral species were identified as those that had <90% RdRp protein identity, or <80% genome identity to previously described viruses. Novel viruses were named using the surnames of figures in the history of Antarctic exploration. Contigs returning blast hits to the RSP13 host reference gene in penguin libraries and the COX1 reference gene in tick libraries were included to compare viral abundance with host marker genes.

To determine whether any viruses identified in ticks were present in the penguin libraries we used Bowtie2 (39) to assemble the raw reads from each penguin library to the novel virus contigs identified in the tick libraries, and vice versa.

### Virus genome annotation and phylogenetic analysis

Viruses were annotated as previously described (32, 33). Viruses with full-length genomes, or incomplete genomes possessing the full-length RNA-dependant RNA polymerase (RdRp) gene, were used for phylogenetic analysis. Amino acid sequences were aligned using MAFFT (40), with poorly aligned sites removed using trimAL (41). The most appropriate model of amino acid substitution was determined for each data set using IQ-TREE (42) or PhyML 3.0 (43), and maximum likelihood (ML) trees were then estimated using PhyML. For initial family and clade level trees, SH-like branch support was used to determine the topological support for individual nodes. Virus clusters providing the most relevant background information to the novel viruses identified in here were then extracted and phylogenetic analysis repeated using PhyML with 1000 bootstrap replicates. In the case of previously described viruses, phylogenies were also estimated using nucleotide sequences.

### Viral diversity and abundance across libraries

Relative virus abundance was estimated as the proportion of the total number of viral reads in each library (excluding rRNA). All ecological measures in the penguin libraries were calculated only using viruses likely associated with birds. Host-association is less complex in tick samples, and in this case we used the full tick data set, only excluding the *Leviviridae* that are associated with bacterial hosts. Analyses were performed using R v 3.4.0 integrated into RStudio v 1.0.143, and plotted using *ggplot2*.

Both the observed virome richness and Shannon effective [alpha] diversity were calculated for each library at the virus family and genus levels using the Rhea script sets (44), and compared between avian orders using the Kruskal-Wallis rank sum test. Beta diversity was calculated using the Bray Curtis dissimilarity matrix and virome structure was plotted as a function of nonmetric multidimensional scaling (NMDS) ordination and tested using Adonis tests using the *vegan* (45) and *phyloseq* packages (46).

## Results

### Diversity and composition of Antarctic penguin and tick viromes

We characterized the transcriptomes of six libraries comprising 10 individuals each, corresponding to three Antarctic penguin species in three locations and four tick libraries comprising a total of 20 ticks (Table S1, Fig 1). RNA sequencing of rRNA depleted libraries resulted in 42,382,642 - 55,930,902 reads assembled into 189,464 - 530,470 contigs for each of the penguin libraries. The tick libraries contained 51,498,136 - 55,930,902 reads assembled into 55,611 - 223,554 contigs (Table S1). There was a large range in the total viral abundance in both the penguin (0.07-0.7 % total viral reads; 0-0.15 % avian viral reads) and tick libraries (0.03-2.4%) (Table S1, Fig 2). In addition to likely avian viruses, the penguin libraries contained numerous reads matching insect, plant, or bacterial viruses and retroviruses (Fig 2A, Fig 3). Retroviruses were excluded from later analyses due to the challenges associated with differentiating exogenous from endogenous sequences using meta-transcriptomic data alone.

**Figure 2.**
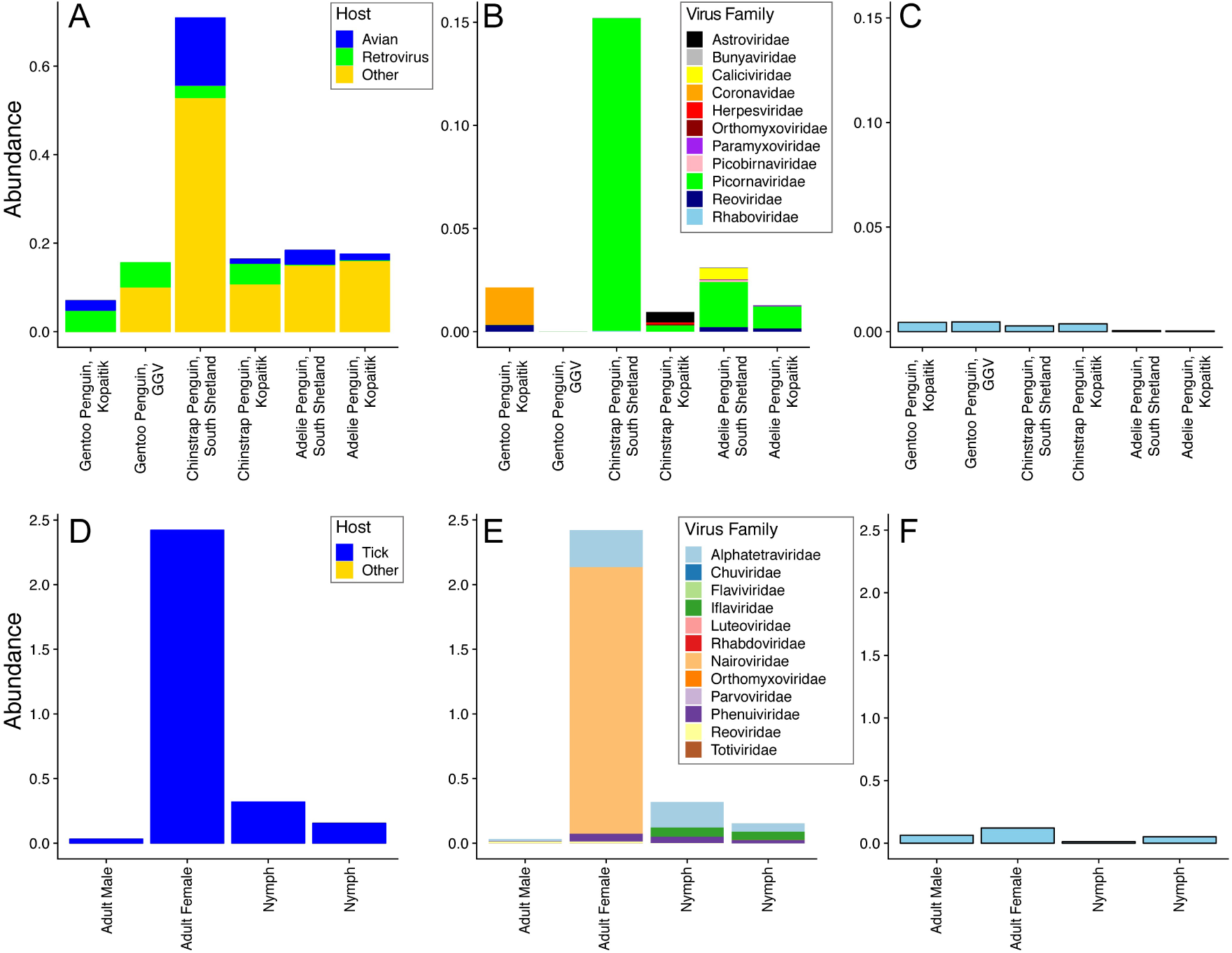
Abundance of viruses found in penguins and their ticks. (A) Abundance of all viral reads found in penguin libraries. (B) Abundance and diversity of avian viruses in each of the penguin libraries. (C) Abundance of the host reference gene RSP13 in penguin libraries. (D) Abundance of all viral reads found in the tick libraries. (E) Abundance and diversity of viruses in each of the tick libraries. (F) Abundance of the host reference gene COX1 in the tick libraries.

**Figure 3.**
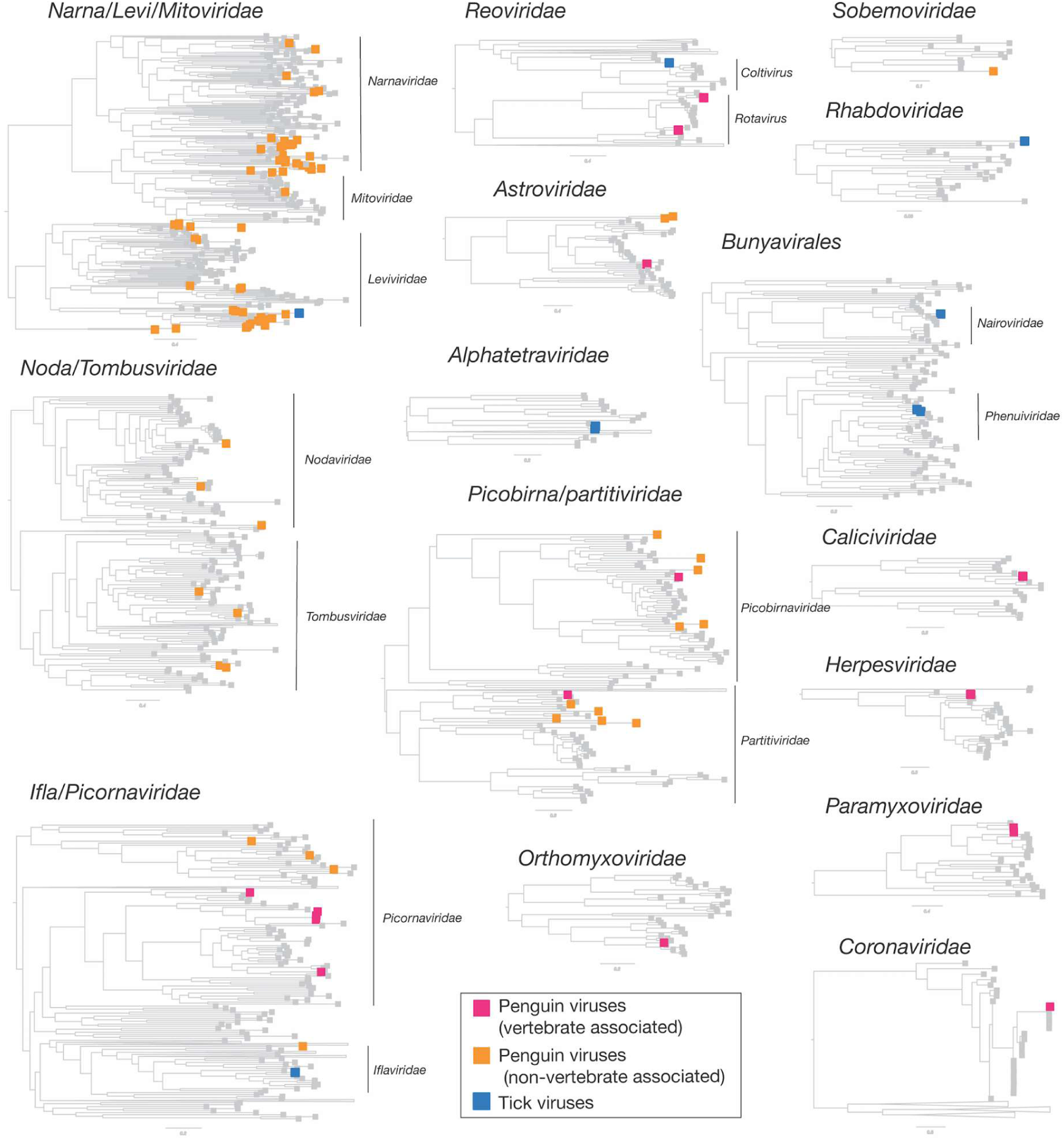
Phylogenetic overview of the viruses found in penguins and ticks. Viruses found in penguins were divided into two groups – those that infect birds and those that likely to other hosts.

The abundance of RSP13, a stably expressed host reference gene in the avian colon (47), was similar across all penguin libraries, yet with lower abundance in the Adelie penguins (Fig 2C). The abundance of COX1 in tick libraries was consistent with the body size of the ticks included in each library, with the highest abundance in the large adult female ticks and the lowest abundance in the first of two nymph libraries (Fig 2D-F).

The abundance of avian viral reads was the highest in Chinstrap penguins sampled on Kopaitik Island (0.152% of total reads), and the lowest in Gentoo penguins sampled at the GGV Base for which no avian viral reads were detected. Because this colony is comprised solely of Gentoo penguins only this species was sampled (31). Alpha diversity (the diversity within each library) was highest in the Adelie and Chinstrap penguins at both the viral family and genus levels, and was lower in Gentoo penguins, even when only considering viromes from Kopaitik Island where all three penguin species were sampled (Fig S1, S2). Hence, the reason we detected no viral reads at the most southern sampling site (the GGV Base) may be due to a combination of location and species choice (Gentoo penguins).

Although there was variation in virus composition among libraries, members of the *Picornaviridae* were the most abundant in the Chinstrap and Adelie penguin libraries, comprising 99%, 32%, 83% and 71% of all the avian viral reads in these four libraries. In marked contrast, the *Picornaviridae* comprised only 0.25% of avian viral reads from Gentoo penguins on Kopaitik Island (and no avian viruses were found in the Gentoo penguins from GGV). Beta diversity demonstrated connectivity in the viromes from the Adelie penguins, driven by a number of shared viral species across the Kopaitik Island and King George Island sampling locations that are 130 km apart (Fig 3, Fig S3).

Within the tick libraries, the greatest virus abundance was seen in the adult female ticks, while the lowest virus abundance was observed in the adult male ticks. Alpha diversity was similar across all libraries. Interestingly, while virus richness was highest in the adult female ticks, Shannon diversity was lower than the other libraries (Fig S4), although without replicates it is not possible to draw clear conclusions. Given the high virus richness in female ticks, it is not surprising that the largest number of viral species were also described in this library. The tick libraries were also highly connected, with 5/8 species shared among them, although the beta diversity calculations are confounded because of limited sample size (Fig S3).

### Substantial RNA virus diversity in Antarctic penguins and their ticks

Overall, 22 viral families, in addition to four viruses that fell outside well defined viral families but clustered with other unclassified ‘picorna-like’ viruses (Treshnikov virus, Luncke virus, Dralkin virus and Tolstivok virus), were identified in the penguin and tick libraries (Fig 3). Of these, the likely bird associated viruses were members of the *Astroviridae, Caliciviridae, Coronaviridae, Herpesviridae, Orthomyxoviridae, Paramyxoviridae, Picorbirnaviridae, Picornaviridae* and *Reoviridae* (Fig 2B, Fig 3) (see below). Ten of the 13 avian associated viruses identified in the penguins likely represent novel avian viral species (Table S2, Fig 4), and two virus species (Avian avulavirus 17 and Shirase virus) were shared among Adelie penguins from different locations. There was no virus sharing among species at individual locations (i.e. no viruses were shared across penguin species at either Kopaitik Island or on King George Island) (Fig 4), although this is likely because species were sampled in different years. All viruses in the tick libraries, with the exception of Taggert virus, represented novel virus species with amino acid sequence similarity to reference virus sequences from 31% to 81%. Five of the nine virus species described in ticks were shared across libraries, which is unsurprising given that the ticks were collected from the same population. Notably, the nymph libraries contained a higher percentage of viral RNA than both the large nymphs and adult male ticks.

**Figure 4.**
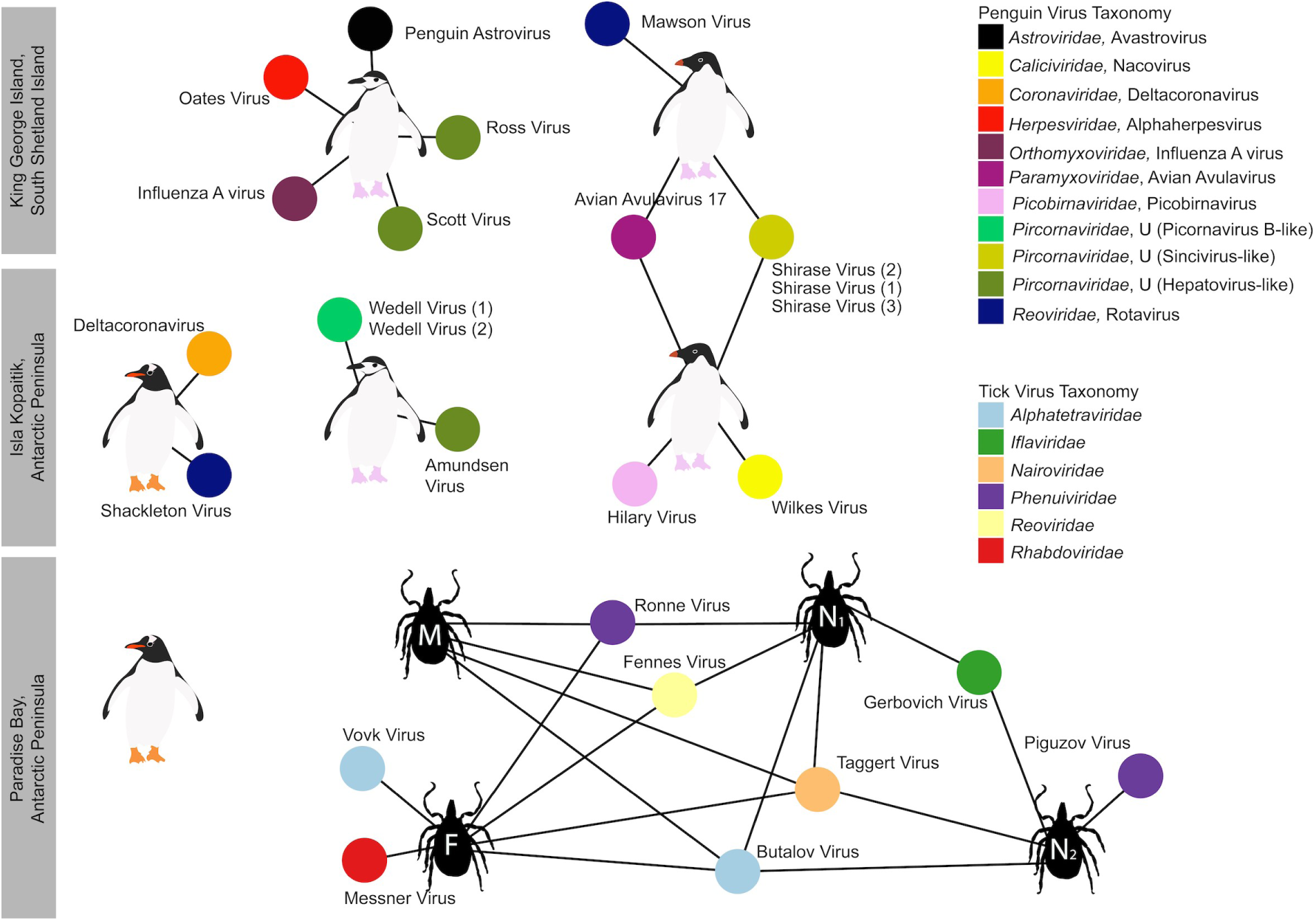
Bipartite network illustrating biologically relevant virus species for which viral genomes were found in each library. Each library is represented as a central node, with a pictogram of the species, surrounded by each viral species. Where two libraries share a virus species, the networks between the two libraries are linked. Virus colour corresponds to virus taxonomy. Viruses identified in penguin libraries that are unlikely to be bird associated are not shown. A list of viruses from each libraries is presented in Table S3, and phylogenetic trees for each virus family can be found in Figs 5-8, S5-S12).

Strikingly, we identified 82 divergent novel virus species in the penguin libraries that clustered phylogenetically within 11 defined families, as well as three viruses that clustered with a group of unclassified viruses. These unclassified viruses likely associated with penguin diet or their microbiome: fish, invertebrates, plants, fungi and bacteria (Fig 2A, Table S3, Fig 3). The largest diversity was found in the “Narna-Levi”, “Noda-Tombus” and “Picorbirna-Pariti” viral groups (36). A number of viruses were highly divergent, including clusters of novel viruses that fell within the *Narnaviridae* and *Leviviridae* (Table S3, Fig 3). Overall, 56 different species of Narna-Levi viruses were identified in Adelie penguins, comprising approximately half of the Narna-like viruses and 21/25 of the *Leviviridae*: these were likely associated with bird diet or microbiome. All invertebrate associated *Picobirnaviridae* were found in Chinstrap penguin libraries, while a single picobirnavirus identified in an Adelie penguin library was most closely related to other bird associated viruses (see below). As these 82 viruses are unlikely to be associated with penguins or their ticks, they are not described further.

### Novel avian viruses

The novel Wilkes virus was identified in an Adelie penguin on Kopaitik Island, and belongs to the genus *Nacovirus* (*Caliciviridae*) - a group dominated by avian viruses (Fig S5). This virus is closely related to Goose calicivirus and caliciviruses sampled from waterbirds in Australia (i.e. Red-necked Avocet and Pink-eared Duck) (32, 48). All the picornaviruses identified in this study likely belong to novel or unassigned genera (Fig S6). Three different variants of Shirase virus were identified in Adelie penguins, two from King George Island and one from Kopaitik Island. Interestingly, Shirase virus falls as a sister lineage to viruses of the genus *Gallivirus*. Similarly, two variants of Wedell virus were identified in Chinstrap penguins. Wedell virus falls in an unassigned lineage of picornaviruses that other avian viruses identified in metagenomic studies (32, 48). Three additional picornaviruses were identified in Chinstrap penguins - Ross virus, Scott virus and Amundsen virus - that fall basal to members of the genus *Tremovirus*.

Rotaviruses were identified in both Gentoo (Shackleton virus) and Adelie (Mawson virus) penguins. Shackleton virus falls as an outgroup to a clade of rotaviruses recently described in wild birds, which are themselves divergent from Rotavirus G virus, while Mawson virus is a sister group to rotavirus D (Fig 5A). Hilary virus, a picobirnavirus, was found in a clade that contains both avian and mammalian hosts. Interestingly, this virus is most closely related to a human picorbirnavirus, albeit with low amino acid similarity and long branch lengths (Fig S7). Although there certainty as to whether these viruses are bacterial rather than vertebrate associated (49), they are retained here for comparative purposes.

**Figure 5.**
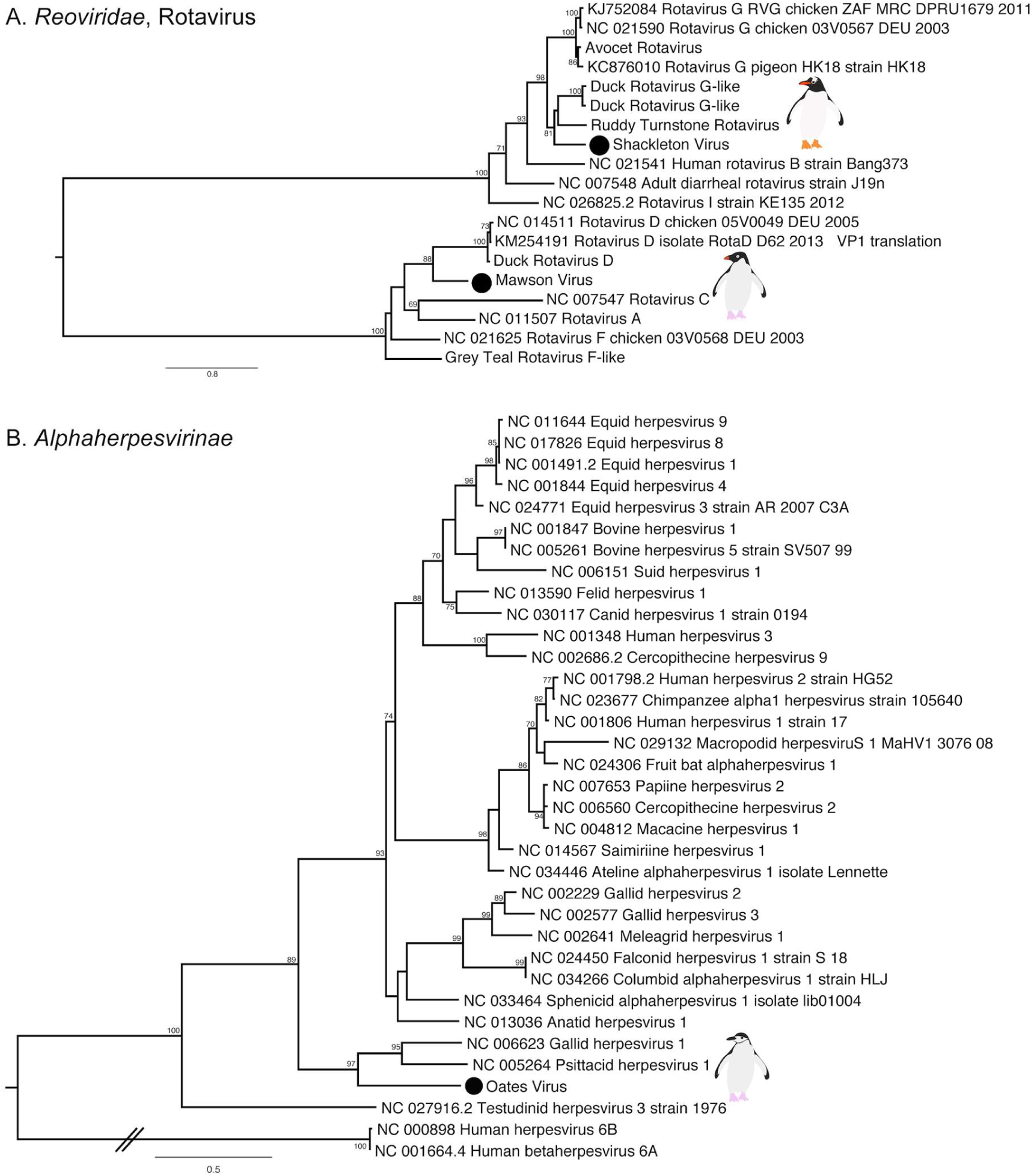
Phylogenies of select novel viruses found in penguins. (A) Phylogenetic tree of the VP1, containing the RdRp, of rotaviruses. The tree is midpoint rooted for clarity only. (B) Phylogeny of the concatenated major capsid gene and glycoprotein B gene of the *Alphaherpesvirinae*. Two betaherpesviruses were used as outgroup to root the tree. The viruses identified in this study are denoted with a filled circle and in bold. Bootstrap values >70% are shown for key nodes. The scale bar represents the number of amino acid substitutions per site.

Finally, although most of the novel viruses documented here had RNA genomes, we also identified a novel alphaherpesvirus, Oates virus, that falls as a sister group to Gallid and Psittacid hepervirus 1. Notably, this virus was distantly related to an alphaherpesvirus previously described in penguins (Sphencid alphaherpesvirus) (Fig 5B).

### Avian RNA viruses previously detected in penguins

Previous studies of Antarctic penguins have detected avian influenza A viruses and avian avulaviruses (15, 16, 21). Similarly, we detected an H5N5 influenza A virus in Chinstrap penguins identical in sequence to that reported previously. This is not surprising as the virus described in Hurt *et al.* (2016) was isolated in the same set of samples (Fig S8). In addition, we identified Avian avulavirus 17 (AAvV-17) in Adelie penguins from both sampling locations (Fig 6A, Fig S9). This virus was previously isolated in Adelie penguins in 2013 (21) and Gentoo penguins in 2014-2016 (50). Analysis of the F gene of AAvV-17 indicates that the virus detected in Adelie penguins on both King George Island and Kopaitik Island was more closely related to that from Gentoo penguins sampled between 2014-2016 (50) than to the Adelie penguins sampled in 2013 (21) (Fig 6A). Although AAvV-17 was detected in penguins sampled at two locations (Kopaitik Island and King George Island) only two weeks apart, they shared only 98.6% identity. Blastx results also indicated the presence of Avian avulavirus 2, although we were unable to assemble the virus genome.

**Figure 6.**
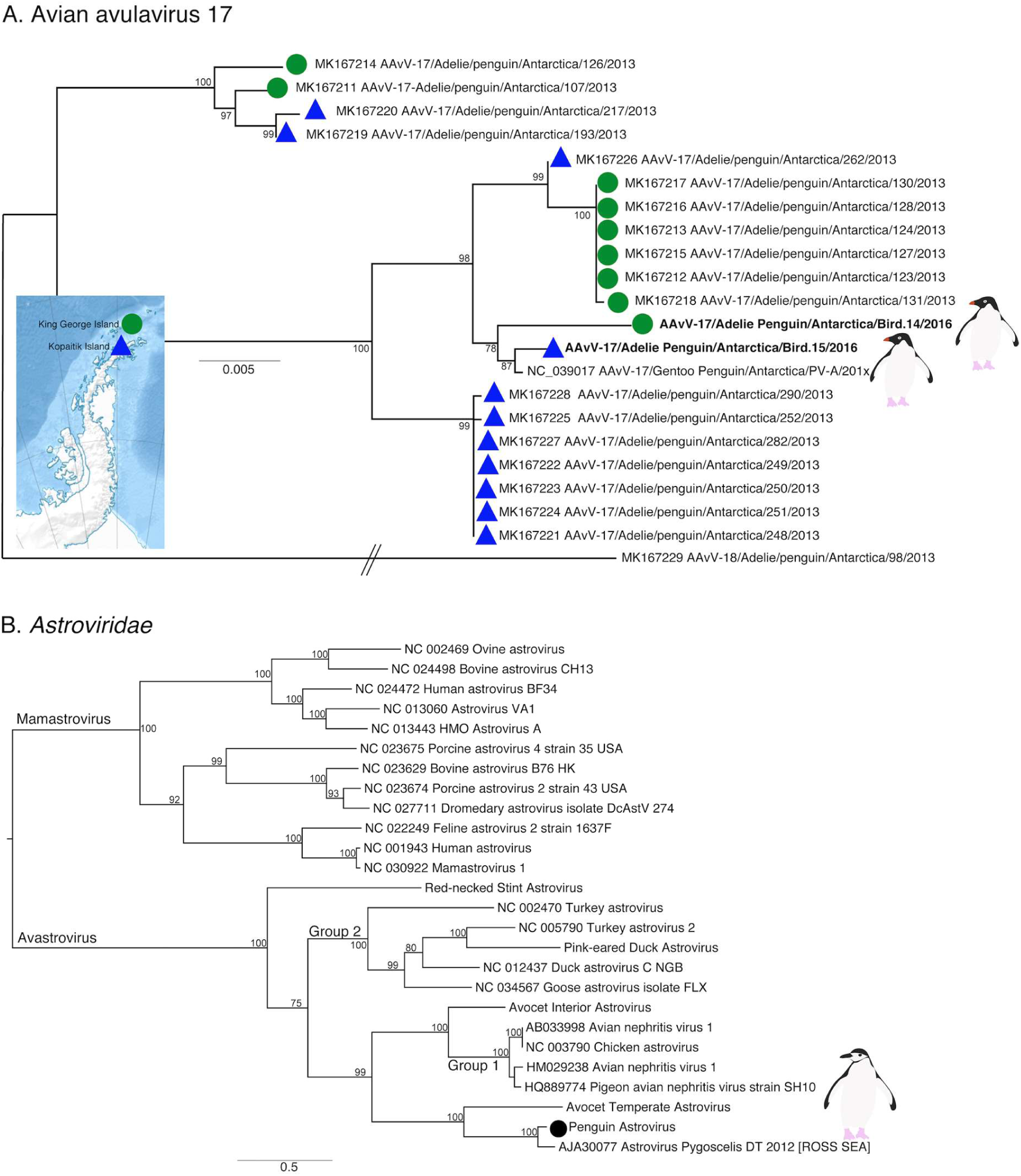
Phylogeny of two previously described viruses in penguins. (A). Phylogeny of the F gene of Avian avulavirus-17. Detection location for viruses identified in this study and Wille *et al.* (2019) are denoted by either a green filled circle (King George Island) or blue filled triangle (Kopaitik Island). It is unclear in which year this virus (AAvV-17/Gentoo Penguin) was isolated (50), although it is sometime between 2014-2016 and therefore has been denoted as 201x. Avian avulavirus 18 was used as outgroup to root the tree. The scale bar represents the number of nucleotide substitutions per site. (B). Phylogenetic tree of the ORF1ab, including the RdRp, of avastroviruses. The tree is mid-point rooted for clarity only. The scale bar represents the number of amino acid substitutions per site. Bootstrap values >70% are shown for key nodes. Viruses identified in this study are denoted in bold.

We also identified a deltacoronavirus and an avastrovirus (Fig 6B, Fig S10-S11). The deltacoronavirus was similar to those reported in birds in the United Arab Emirates, Australia, Niger, and Finland, with ~95% identity. A lack of sampling makes it challenging to determine how deltacoronaviruses in Antarctica and other continents may be shared (Fig S10). The astrovirus detected was similar (88.3% identity) to a short fragment (1000 bp) previously reported in Adelie penguins on the Ross ice shelf of Antarctica (22) (Table S2), a pattern confirmed by phylogenetic analysis (Fig 6B). Phylogenetic analysis also reveals that this virus falls in an outgroup to Group 2 viruses, including Avian Nephritis virus (Fig 6B, Fig S11). Although we were unable to determine the epidemiology of these viruses in Antarctica, repeated detection on opposite ends of the Antarctic continent makes it possible that this is a penguin specific virus.

### Tick associated viruses

The most abundant virus identified within the *I. uriae* ticks sampled here was a variant of Taggert virus, a nairovirus (order *Bunyavirales*) previously identified in penguin associated ticks on Macquarie Island: the contigs identified in our data showed 81.6% nucleotide sequence similarity in the RdRp region to Taggert virus (24) (Fig 7). This Taggert virus variant accounted for 2.0% of total reads (87% of viral reads) in the adult female library and was found in all tick libraries. Because the nucleotide sequences of Taggert virus differed between libraries it is unlikely that they represent cross-library contamination. In addition, we identified 75 reads of Taggert virus in the library containing samples from Chinstrap penguins on Kopaitik Island. Importantly, the tick and penguin libraries were not sequenced on the same lane, or even in the same time frame, thereby excluding contamination. Two other members of the order *Bunyavirales* were also discovered - Ronne virus and Barre virus - both members of the *Phenuiviridae* (Fig S12) that exhibited 80% amino acid sequence similarity across the RdRp segment. Ronne virus was identified in three of the tick libraries (adult male, adult female and nymph library1) but Barre virus was identified only in a single library (nymph library 2) (Fig 4).

**Figure 7.**
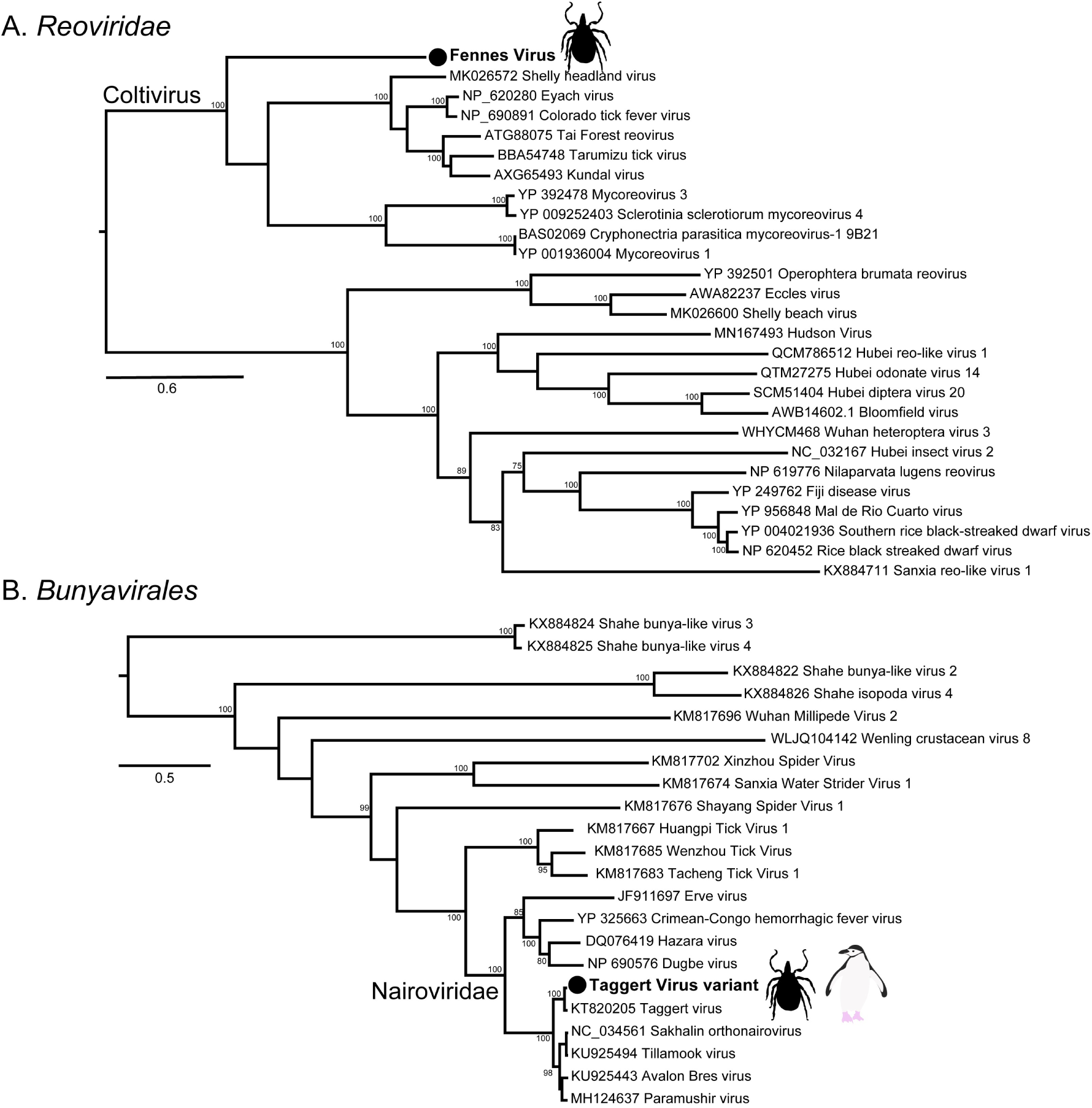
Phylogenies of tick arboviruses. (A) The RdRp segment of select members of the *Reoviridae*, including the genus *Coltivirus*. (B). The RdRp of select members of the *Bunyavirales* including the family *Nairoviridae*. The novel tick viruses identified in this study are denoted with a filled circle and in bold. The tree has been mid-point rooted for clarity only. Bootstrap values >70% are shown for key nodes. The scale bar represents the number of amino acid substitutions per site.

The six other novel virus species identified in the tick libraries comprised five viral families: *Iflaviridae-like, Alphatetraviridae, Reoviridae, Rhabdoviridae*, and *Levivirdae*. A novel Ifla-like virus, Gerbovich virus, was identified within both nymph libraries. This virus clustered with a group of tick associated ifla-like viruses, including Ixodes holocyclus iflavirus and Ixodes scapularis iflavirus (Fig S12). Two sister species of virus were identified within the *Alphatetraviridae* - Bulatov virus and Vovk virus - that showed 76.1% amino acid sequence similarity across the RdRp region. These two viruses are highly divergent from all other RdRp sequences currently available, exhibiting just 35.7% amino acid sequence similarity to the divergent tick-borne tetravirus-like virus (Fig S12). A novel colti-like virus (*Reoviridae*), Fennes virus, was identified in the adult male, female and nymph libraries, although we were only able to assemble four segments. Notably, Fennes virus falls basal to the existing coltivirus group, exhibiting just 30% amino acid sequence similarity to Shelly headland virus, recently identified in *Ixodes holocyclus* ticks from Australia (Fig 7). The partial genome of a *Rhabdovirus*, Messner virus, was identified in the adult female library. However, this fragment was of low abundance, and only the RdRp segment (Fig S12).

Finally, Mackintosh virus, identified in all four tick libraries, was not associated with any other tick viruses. Instead, this virus clustered with viruses from the *Leviviridae* indicating that it is likely a bacteriophage (Table S2).

## Discussion

The advent of metagenomic sequencing and improved sampling has rapidly accelerated the rate of microbial discovery in the Antarctic. Indeed, viruses have now been identified both in the environment (e.g. Antarctic lakes), and in wildlife. We aimed to test the hypothesis that Antarctic penguin colonies experience low pathogen pressure as a result of their geographic and climatic isolation, employing meta-transcriptomic virus discovery from three penguin species and their ticks. Critically, we demonstrate the presence of 13 viral species in these penguins and nine in ticks associated with penguin nesting sites. These data counter the idea that animals in the Antarctic harbour less microbial diversity than animals from other geographic regions. Indeed, the penguins sampled show similar levels of virome diversity as Australian wild birds (32, 48). Recent virome studies of Australian birds revealed an alpha diversity (observed richness) of 5.37 and 5.8 per library, with an average of 2.87 and 3.1 viral genomes and 60% and 80% of viruses being novel (32, 48). In comparison, in the penguins studied here we observed an average richness of 4.6 and 2.8 viral genomes per library, with a virus discovery rate of 76%. There was also an impressive level of viral diversity in the tick libraries considering the small sample size: eight novel virus species and a single previously identified species were identified in 20 ticks, compared to 19 novel viruses in 146 ticks from Australia (33). Finally, in both the penguin and tick associated viruses revealed we identified similar viral families to those documented previously (32, 33, 48). This strongly suggests that these families and genera are associated with a huge diversity of birds and ticks across the globe, providing a viral connectivity between geographically distinct localities.

Notably, as all the penguins sampled appeared healthy, the disease-causing capacity of these viruses is uncertain. Ten Chinstrap penguins sampled on King George Island harboured five different viral species, mostly from the *Picornaviridae*, at very high abundance (0.15% of total reads). Adelie penguins also had high viral diversity, with apparently healthy birds carrying three or four viral species. Perhaps more striking was that Adelie penguins on King George Island and the Antarctic peninsula shared viruses, despite the greater than 100 km distance between these colonies. A similar trend was observed by Wille *et al.* (2019a) who found that Adelie penguins sampled in 2013 shared avian avulavirus 17 and 19 across these two locations, thereby revealing a connectivity between penguin colonies. Whether this is due to overlapping foraging grounds, prospecting birds visiting different colonies, viral vectors in the form of predatory and scavenging birds such as Southern Giant Petrels (*Macronectes gigantes*), Kelp Gulls (*Larus dominicanus*) or Skuas (*Stercorarius spp.*) (51, 52), or another unimagined route is unclear. Penguins sampled on King George Island and Kopaitik Island had similar alpha diversity at the virus family and genus levels. Interestingly, no avian viral reads or genomes were detected in the samples from Gentoo penguins at GGV. Whether this is due to geographic structuring of avian viruses in Antarctica, the species sampled at this location (i.e. Gentoo penguins tended to have lower diversity than either Adelie or Chinstrap penguins) or another process merits further investigation.

Combined, these data strongly suggest that penguins are not merely spill-over hosts, but may be central reservoir hosts for a diverse range of viruses. This is apparent in two observations. The first is the repeated detection of specific virus species, such as avian avulaviruses, and that these viruses comprise distinct clusters of related variants. Antarctic penguins have been sampled since the 1970’s, and avian avulaviruses have repeatedly been detected, both by serology and PCR. The detection of phylogenetically related avian avulavirus 17 in 2013 in Adelie penguins (21), in 2014-2016 in Gentoo penguins (50), and again in 2016 in Adelie penguins as shown here, strongly suggests that these animals are an important reservoir for these viruses. Although the influenza A virus we detected was the same virus as described previously (15), the long branch lengths in the phylogenetic trees suggest long-term undetected circulation in Antarctica (15).

The second key observation that indicates that penguins are potential virus reservoirs was the presence of likely arboviruses, which is why we paired our analysis of the penguin virome with that of the ticks that parasitise them. Of the nine species of viruses identified within the ticks in this study, two clustered phylogenetically with other arboviruses: the previously detected Taggert virus fell within the *Nairoviridae*, while Fennes virus was a member of the *Reoviridae*. Taggert virus was originally identified in *I. uriae* collected from penguin colonies, and is one of seven *I. uriae* associated virus species identified in ticks collected from penguin colonies on Macquarie Island (24). Taggert virus groups phylogenetically within the genus *Orthonairovirus*, and closely to the pathogenic arbovirus, Crimean-Congo Haemorrhagic fever virus. Interestingly, we not only identified Taggert virus in all four tick libraries, but also in Chinstrap penguins on Kopaitik Island. This strongly supports the idea that penguins acted as a reservoir host for Taggert virus (24). In this context it is important to note that as all penguin and tick samples were processed in separate laboratories and sequenced separately, thereby eliminating cross-library contamination.

Also of note was Fennes virus that clustered phylogenetically within the genus *Coltivirus* that includes the pathogenic tick-borne virus Colorado tick fever virus as well as a number of tick associated viruses and a species identified in African bats (33, 53–55). Notably, Fennes virus fell in a basal position and was relatively divergent from the other coltiviruses. The vertebrate reservoirs of coltiviruses have been only confirmed for Colorado tick fever virus and Tai forest reovirus - rodents and free tailed bats, respectively - although other members of the genus are suspected to infect rodent species. Interestingly, the viruses identified in *I. uriae* from Macquarie Island in a series of three studies between the 1970s and 2009 belonged to just four families - *Reoviridae*, *Narioviridae*, *Phenuiviridae* and *Flaviviridae* (23–25) - three of which were present here. Our phylogenetic analysis suggested that six of the remaining seven virus species identified within the tick data were likely associated with invertebrates, with one other virus (Mackintosh virus) likely a bacteriophage from the family *Leviviridae*.

There was extensive diversity of viruses identified in the penguin samples that were likely to be associated with hosts other than birds, including entire clades of novel viruses within phylogenetic trees of the *Narnaviridae* and *Leviviridae*. Given their phylogenetic position these viruses are likely associated with the fish, crustacean and plant species ingested by the penguins as part of their diet, as well as infecting unicellular parasites, fungi and bacteria. A number of these viruses may also be associated with penguin gut flora: indeed, cloacal swabs are used extensively in studies of bird gut microbiomes (56, 57). Due to the nature of the cloacal swabs, it is impossible to accurately determine the host for these viruses, although some information can be gleaned from the families in which these virus fall. For example, the *Narnaviridae* are known to infect fungi and protists, while the related *Leviviridae* infect bacteria (58–61). Other novel viruses fell within invertebrate associated clades of the *Nodaviridae* and *Tombusviridae*, associated with both vertebrate and invertebrate infecting viruses in the *Picornavirales* (36). There were also a number of novel viruses identified that clustered within the *Picobirnaviridae/Partitiviridae* group. While the *Partitiviridae* are recognised as invertebrate associated viruses, the host association of the *Picobirnaviridae* is currently uncertain (49). Overall, this demonstrates remarkable undescribed viral richness in those organisms that comprise the diet of Antarctic penguins.

In sum, we reveal substantial viral diversity in Antarctic penguins, their diet and their ticks. We therefore expect that additional viruses will be identified with increased sampling, reflecting what it is in reality a relatively high diversity of unique fauna and flora on the Antarctic continent. Clearly, additional sampling of penguins and other species in Antarctica is critical to elucidate the epidemiological connection between Antarctica and the rest of the globe, and from this better understand the mechanisms of viral introduction and circulation.

## Acknowledgements

The Melbourne WHO Collaborating Centre for Reference and Research on Influenza is supported by the Australian Department for Health. ECH is funded by an ARC Australian Laureate Fellowship (FL170100022). The fieldwork was funded by the Instituto Antártico Chileno as part of the project INACH T-27-INACH T-12-13:” *Campylobacter* in Antarctica: diversity, origin and effects on wildlife”. We are grateful to Jorge Hernández, Gonzalo Medina, Lucila Moreno, Juana López, Patrik Ellström, Michelle Thomson, Fernanda González, Håkan Johansson for assistance in field collection, and the staff of the Chilean bases for their hospitality.

## References

1. Grimaldi WW, Seddon PJ, Lyver PO, Nakagawa S, Tompkins DM. Infectious diseases of Antarctic penguins: current status and future threats. Polar Biol. 2015;38:591–606.

2. Dobson A, Foufopoulos J. Emerging infectious pathogens of wildlife. Philos T R Soc B. 2001;356:1001–12.

3. Sutherland WJ, Aveling R, Bennun L, Chapman E, Clout M, Cote IM, et al. A horizon scan of global conservation issues for 2012. Trends Ecol Evol. 2012;27:12–8.

4. Hughes KA, Convey P. The protection of Antarctic terrestrial ecosystems from inter- and intra-continental transfer of non-indigenous species by human activities: A review of current systems and practices. Global Environ Chang. 2010;20:96–112.

5. Chown SL, Lee JE, Hughes KA, Barnes J, Barrett PJ, Bergstrom DM, et al. Challenges to the Future Conservation of the Antarctic. Science. 2012;337:158–9.

6. Smeele ZE, Ainley DG, Varsani A. Viruses associated with Antarctic wildlife: From serology based detection to identification of genomes using high throughput sequencing. Virus Res. 2018;243:91–105.

7. Morgan IR, Westbury HA. Virological studies of Adelie Penguins (Pygoscelis adelia) in Antarctica. Avian Dis. 1981;25:1019–26.

8. Morgan IR, Westbury HA, Campbell J. Viral infections of Little Blue Penguins (Eudyptula minor) along the southern coast of Australia. J Wildlife Dis. 1985;21:193–8.

9. Gardner H, Kerry K, Riddle M, Brouwer S, Gleeson L. Poultry virus infection in Antarctic penguins. Nature. 1997;387:245-.

10. Lang AS, Lebarbenchon C, Ramey AM, Robertson GJ, Waldenström J, Wille M. Assessing the role of seabirds in the ecology of influenza A viruses. Avian Dis. 2016;60:378–86.

11. Wallensten A, Munster VJ, Osterhaus ADME, Waldenström J, Bonnedahl J, Broman T, et al. Mounting evidence for the presence of influenza A virus in the avifauna of the Antarctic region. Antarct Sci. 2006;18.

12. Austin FJ, Webster RG. Evidence of ortho- and paramyxoviruses in fauna from Antarctica. J Wildl Dis. 1993;29:568–71.

13. Grimaldi W, Ainley DG, Massaro M. Multi-year serological evaluation of three viral agents in the Adelie Penguin (Pygoscelis adeliae) on Ross Island, Antarctica. Polar Biol. 2018;41:2023–31.

14. Thomazelli LM, Araujo J, Oliveira DB, Sanfilippo L, Ferreira CS, Brentano L, et al. Newcastle disease virus in penguins from King George Island on the Antarctic region. Vet Microbiol. 2010;146:155–60.

15. Hurt AC, Su YC, Aban M, Peck H, Lau H, Baas C, et al. Evidence for the introduction, reassortment, and persistence of diverse influenza A viruses in Antarctica. J Virol. 2016;90:9674–82.

16. Hurt AC, Vijaykrishna D, Butler J, Baas C, Maurer-Stroh S, Silva-de-la-Fuente MC, et al. Detection of evolutionarily distinct avian influenza A viruses in Antarctica. MBio. 2014;5:e01098–14. doi: 10.1128/mBio.-14.

17. Lee SY, Kim JH, Park YM, Shin OS, Kim H, Choi HG, et al. A novel adenovirus in Chinstrap Penguins (Pygoscelis antarctica) in Antarctica. Viruses. 2014;6:2052–61.

18. Lee SY, Kim JH, Seo TK, No JS, Kim H, Kim WK, et al. Genetic and molecular Epidemiological characterization of a novel Adenovirus in Antarctic penguins collected between 2008 and 2013. PLoS ONE. 2016;11.

19. Varsani A, Kraberger S, Jennings S, Porzig EL, Julian L, Massaro M, et al. A novel papillomavirus in Adelie penguin (Pygoscelis adeliae) faeces sampled at the Cape Crozier colony, Antarctica. J Gen Virol. 2014;95:1352–65.

20. Varsani A, Porzig EL, Jennings S, Kraberger S, Farkas K, Julian L, et al. Identification of an avian polyomavirus associated with Adelie penguins (Pygoscelis adeliae). J Gen Virol. 2015;96:851–7.

21. Wille M, Aban M, Wang J, Moore N, Shan S, Marshall J, et al. Antarctic penguins as reservoirs of diversity for avian avulaviruses. J Virol. 2019 doi: 10.1128/JVI.00271-19.

22. Grimaldi WW, Hall RJ, White DD, Wang J, Massaro M, Tompkins DM. First report of a feather loss condition in Adelie penguins (Pygoscelis adeliae) on Ross Island, Antarctica, and a preliminary investigation of its cause. Emu. 2015;115:185–9.

23. Major L, La Linn M, Slade RW, Schroder WA, Hyatt AD, Gardner J, et al. Ticks associated with Macquarie Island penguins carry arboviruses from four genera. PLoS ONE. 2009;4.

24. Doherty RL, Carley JG, Murray MD, Main AJ, Kay BH, Domrow R. Isolation of arboviruses (Kemerovo-Group, Sakhalin-Group) from Ixodes uriae collected at Macquarie Island, Southern Ocean. Am J Trop Med Hyg. 1975;24:521–6.

25. Stgeorge TD, Doherty RL, Carley JG, Filippich C, Brescia A, Casals J, et al. The isolation of arboviruses including a new Flavivirus and a new Bunyavirus from Ixodes-(Ceratixodes)-uriae (Ixodoidea, Ixodidae) collected at Macquarie Island, Australia, 1975-1979. Am J Trop Med Hyg. 1985;34:406–12.

26. Barbosa A, Benzal J, Vidal V, D’Amico V, Coria N, Diaz J, et al. Seabird ticks (Ixodes uriae) distribution along the Antarctic Peninsula. Polar Biol. 2011;34:1621–4.

27. Munoz-Leal S, Gonzalez-Acuna D. The tick Ixodes uriae (Acari: Ixodidae): Hosts, geographical distribution, and vector roles. Ticks Tick Borne Dis. 2015;6:843–68.

28. Benoit JB, Lopez-Martinez G, Elnitsky MA, Lee RE, Denlinger DL. Increase in feeding by the tick, Ixodes uriae, on Adelie penguins during a prolonged summer. Antarct Sci. 2009;21:151–2.

29. Benoit JB, Yoder JA, Lopez-Martinez G, Elnitsky MA, Lee RE, Denlinger DL. Habitat requirements of the seabird tick, Ixodes uriae (Acari: Ixodidae), from the Antarctic Peninsula in relation to water balance characteristics of eggs, nonfed and engorged stages. J Comp Physiol B. 2007;177:205–15.

30. Frenot Y, de Oliveira E, Gauthier-Clerc M, Deunff J, Bellido A, Vernon P. Life cycle of the tick Ixodes uriae in penguin colonies: relationships with host breeding activity. Int J Parasitol. 2001;31:1040–7.

31. Humphries GRW, Che-Castaldo C, Naveen R, Schwaller M, McDowall P, Schrimpf M, et al. Mapping Application for Penguin Populations and Projected Dynamics (MAPPPD): Data and tools for dynamic management and decision support. Polar Rec. 2017 www.penguinmap.com/mappd..

32. Wille M, Eden JS, Shi M, Klaassen M, Hurt AC, Holmes EC. Virus-virus interactions and host ecology are associated with RNA virome structure in wild birds. Mol Ecol. 2018 doi: 10.1111/mec.14918.

33. Harvey E, Rose K, Eden JS, Lo N, Abeyasuriya T, Shi M, et al. Extensive diversity of RNA viruses in Australian ticks. J Virol. 2019; 93.

34. Grabherr MG, Haas BJ, Yassour M, Levin JZ, Thompson DA, Amit I, et al. Full-length transcriptome assembly from RNA-Seq data without a reference genome. Nature Biotech. 2011;29:644–52.

35. Buchfink B, Xie C, Huson DH. Fast and sensitive protein alignment using DIAMOND. Nat Methods. 2015;12:59–60.

36. Shi M, Lin XD, Tian JH, Chen LJ, Chen X, Li CX, et al. Redefining the invertebrate RNA virosphere. Nature. 2016;540:539–43.

37. Shi M, Lin XD, Chen X, Tian JH, Chen LJ, Li K, et al. The evolutionary history of vertebrate RNA viruses. Nature. 2018;556:197–202.

38. Asplund M, Kjartansdottir KR, Mollerup S, Vinner L, Fridholm H, Herrera JAR, et al. Contaminating viral sequences in high-throughput sequencing viromics: a linkage study of 700 sequencing libraries. Clin Microbiol Infect. 2019.

39. Langmead B, Salzberg SL. Fast gapped-read alignment with Bowtie 2. Nature Methods. 2012;9:357–U54.

40. Katoh K, Standley DM. MAFFT multiple sequence alignment software version 7: improvements in performance and usability. Mol Biol Evol. 2013;30:772–80.

41. Capella-Gutierrez S, Silla-Martinez JM, Gabaldon T. trimAl: a tool for automated alignment trimming in large-scale phylogenetic analyses. Bioinformatics. 2009;25:1972–3.

42. Nguyen LT, Schmidt HA, von Haeseler A, Minh BQ. IQ-TREE: A Fast and Effective Stochastic Algorithm for Estimating Maximum-Likelihood Phylogenies. Mol Biol Evol. 2015;32:268–74.

43. Guindon S, Dufayard JF, Lefort V, Anisimova M, Hordijk W, Gascuel O. New algorithms and methods to estimate maximum-likelihood phylogenies: assessing the performance of PhyML 3.0. Syst Biol. 2010;59:307–21.

44. Lagkouvardos I, Fischer S, Kumar N, Clavel T. Rhea: a transparent and modular R pipeline for microbial profiling based on 16S rRNA gene amplicons. PeerJ. 2017;5:e2836. doi: 10.7717/peerj.2836.

45. Oksanen J, Kindt R, Legendre P, O’Hara B, Stevens MHH, Oksanen MJ, et al. The vegan package. Commun Ecol Package. 2007;10:631–7.

46. McMurdie PJ, Holmes S. phyloseq: An R package for reproducible interactive analysis and graphics of microbiome census data. PLoS ONE. 2013;8:e61217. doi: 10.1371/journal.pone.0061217.

47. Chapman JR, Helin AS, Wille M, Atterby C, Jarhult JD, Fridlund JS, et al. A panel of stably expressed reference genes for real-time qPCR gene expression studies of Mallards (Anas platyrhynchos). PLoS ONE. 2016;11:e0149454. doi: 10.1371/journal.pone.

48. Wille M, Shi M, Klaassen M, Hurt AC, Holmes EC. Virome heterogeneity and connectivity in waterfowl and shorebird communities. ISME J. 2019;13:2603–16.

49. Krishnamurthy SR, Wang D. Extensive conservation of prokaryotic ribosomal binding sites in known and novel picobirnaviruses. Virology. 2018;516:108–14.

50. Neira V, Tapia R, Verdugo C, Barriga G, Mor S, Ng TFF, et al. Novel avulaviruses in penguins, Antarctica. Emerg Infect Dis. 2017;23:1212–4.

51. Leotta GA, Chinen I, Vigo GB, Pecoraro M, Rivas M. Outbreaks of avian cholera in Hope Bay, Antarctica. J Wildlife Dis. 2006;42:259–70.

52. Baumeister E, Leotta G, Pontoriero A, Campos A, Montalti D, Vigo G, et al. Serological evidences of influenza A virus infection in Antarctica migratory birds. Int Congr. 2004;1263:737–40.

53. Weiss S, Dabrowski PW, Kurth A, Leendertz SAJ, Leendertz FH. A novel Coltivirus-related virus isolated from free-tailed bats from Cote d’Ivoire is able to infect human cells in vitro. Virol J. 2017;14.

54. Fujita R, Ejiri H, Lim CK, Noda S, Yamauchi T, Watanabe M, et al. Isolation and characterization of Tarumizu tick virus: A new coltivirus from Haemaphysalis flava ticks in Japan. Virus Res. 2017;242:131–40.

55. Goodpasture HC, Poland JD, Francy B, Bowen GS, Horn KA. Colorado tick fever - clinical, epidemiologic, and laboratory aspects of 228 cases in Colorado in 1973-1974. Ann Intern Med. 1978;88:303–10.

56. Risely A, Waite D, Ujvari B, Hoye B, Klaassen M. Active migration is associated with specific and consistent changes to gut microbiota in Calidris shorebirds. J Animal Ecol. 2017 doi: 10.1111/365-2656.12784.

57. Risely A, Waite D, Ujvari B, Klaassen M, Hoye B. Gut microbiota of a long-distance migrant demonstrates resistance against environmental microbe incursions. Mol Ecol. 2017;26:5842–54.

58. Hillman BI, Cai G. The family Narnaviridae: simplest of RNA viruses. Adv Virus Res. 2013;86:149–76.

59. Grybchuk D, Kostygov AY, Macedo DH, Votypka J, Lukes J, Yurchenko V. RNA Viruses in Blechomonas (Trypanosomatidae) and evolution of Leishmaniavirus. MBio. 2018;9.

60. Akopyants NS, Lye LF, Dobson DE, Lukes J, Beverley SM. A narnavirus in the trypanosomatid protist plant pathogen Phytomonas serpens. Genome Announc. 2016;4.

61. Callanan J, Stockdale SR, Shkoporov A, Draper LA, Ross RP, Hill C. RNA phage biology in a metagenomic era. Viruses. 2018;10.

